# Neutralizing activity of broadly neutralizing anti-HIV-1 antibodies against primary African isolates

**DOI:** 10.1101/2020.09.24.310938

**Authors:** Julio C.C. Lorenzi, Pilar Mendoza, Yehuda Z. Cohen, Lilian Nogueira, Christy Lavine, Joseph Sapiente, Michael S. Seaman, Marie Wiatr, Nelly R. Mugo, Andrew Mujugira, Sinead Delany, Jairam Lingappa, Connie Celum, Marina Caskey, Michel C. Nussenzweig

**Affiliations:** Laboratory of Molecular Immunology, The Rockefeller University, New York, NY, USA; Center for Virology and Vaccine Research, Beth Israel Deaconess Medical Center, Harvard Medical School, Boston, MA, USA; Kenya Medical Research Institute (KEMRI), Nairobi, Kenya; Infectious Diseases Institute, Makerere University, Kampala, Uganda; University of the Witswatersrand, South Africa; Department of Global Health, University of Washington, Seattle, Washington, USA; Department of Medicine, University of Washington, Seattle, Washington, USA; Department of Pediatrics, University of Washington, Seattle, Washington, USA; Department of Epidemiology, University of Washington, Seattle, Washington, USA; Howard Hughes Medical Institute

## Abstract

Novel therapeutic and preventive strategies are needed to contain the HIV-1 epidemic. Broadly neutralizing human antibodies (bNAbs) with exceptional activity against HIV-1 are currently being tested in HIV-1 prevention trials. The selection of anti-HIV-1 bNAbs for clinical development was primarily guided by their *in vitro* neutralizing activity against HIV-1 Env pseudotyped viruses. Here we report on the neutralizing activity of 9 anti-HIV-1 bNAbs now in clinical development against 126 Clade A, C, D PBMC-derived primary African isolates. The neutralizing potency and breadth of the bNAbs tested was significantly reduced compared to pseudotyped viruses panels. The difference in sensitivity between pseudotyped viruses and primary isolates varied from 3-to nearly 100-fold depending on the bNAb and the HIV-1 clade. Thus, the neutralizing activity of bNAbs against primary African isolates differs and cannot be predicted from their activity against pseudovirus panels. The data have significant implications for interpreting the results of ongoing HIV-1 prevention trials.

## Introduction

The development of broadly neutralizing antibodies (bNAbs) against HIV has matured in the past few years as several bNAbs were evaluated in clinical trials. VRC01 (Wu et al., 2010), 3BNC117 (Scheid et al., 2011) and 10-1074 (Mouquet et al., 2012) have been the most extensively evaluated to date, in healthy volunteers (Cohen et al., 2019; Mayer et al., 2017), viremic people living with HIV (Caskey et al., 2015, 2017; Lynch et al., 2015; Bar-On et al., 2018), and in the setting of analytical treatment interruption (Scheid et al., 2016; Cohen et al., 2018a; Mendoza et al., 2018; Bar et al., 2016), with encouraging results. These trials were restricted to patients living in the United States and Europe, limiting the assessment of the global utility of these antibodies, since the majority of the individuals in the regions in question were infected with clade B HIV-1 (Bbosa et al., 2019).

One potential limitation in the development of bNAbs is that their activity has been documented primarily using panels of Env-pseudotyped viruses. However, we (Cohen et al., 2018b) and others (Louder et al., 2005; Mann et al., 2009; Provine et al., 2012; Etemad et al., 2015) have shown that Env-pseudotyped viruses often overestimate both the breadth and potency of bNAbs as compared to PBMC-derived HIV isolates.

Here we report on the breadth and potency of nine bNAbs currently in clinical development against primary PBMC-derived HIV-1 viruses isolated from individuals living in South Africa, Uganda, and Kenya. We compared these results with data from Env-pseudotyped virus panels as well as matched Env-pseudotyped viruses derived from the African isolates.

## Results

To examine the coverage of bNAbs in clinical development against HIV-1 variants circulating in Africa, we obtained 218 cryopreserved PBMC samples from people living with HIV-1 who participated in one of three studies: The Partners in Prevention HSV/HIV Transmission Study (Celum et al., 2010), The Couples Observational Study (Lingappa et al., 2011), or The Partners PrEP Study (Baeten et al., 2012). The samples were collected from participants recruited at sites in South Africa (N=84), Uganda (N=68), and Kenya (N=66). Bulk CD4^+^ T lymphocytes were cultured yielding 126 (58%) HIV-1 isolates after 21 days (Supp. Table 1).

**Table 1.**
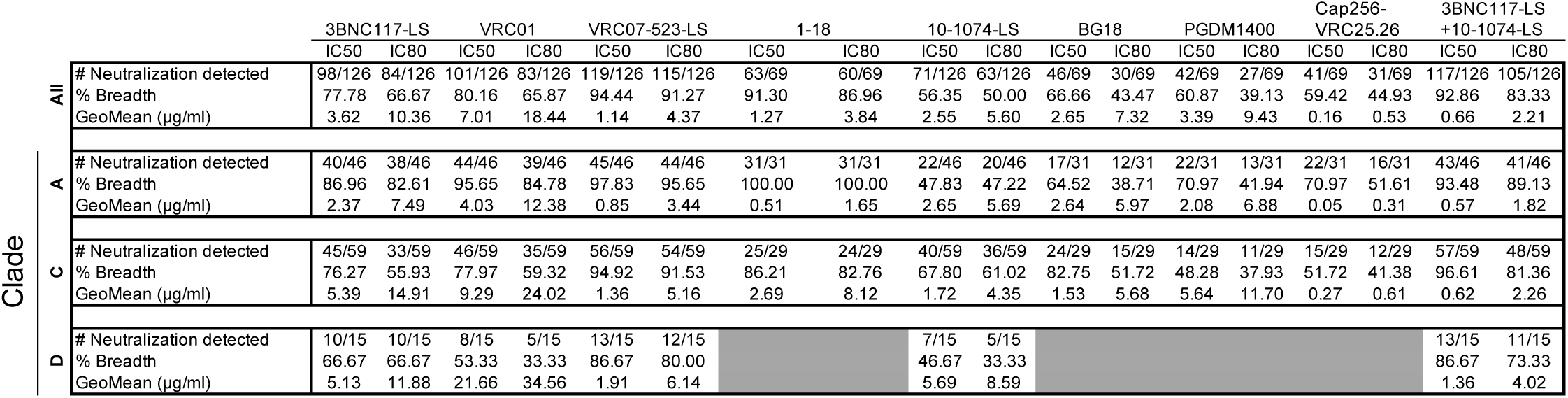
Breadth, IC_50_s _and_ IC_80_s in TZM-bl cells for PBMC derived isolates

To examine the genetic diversity of the HIV-1 viruses obtained from the cultures, we performed single genome amplification (SGA) on 53 viral supernatants and obtained 172 independent sequences representing clades A, C, and D, with 2 sequences per supernatant on average. We observed that the viruses were phylogenetically grouped in large part by their geographic origins and clades (Fig. 1a).

**Figure 1.**
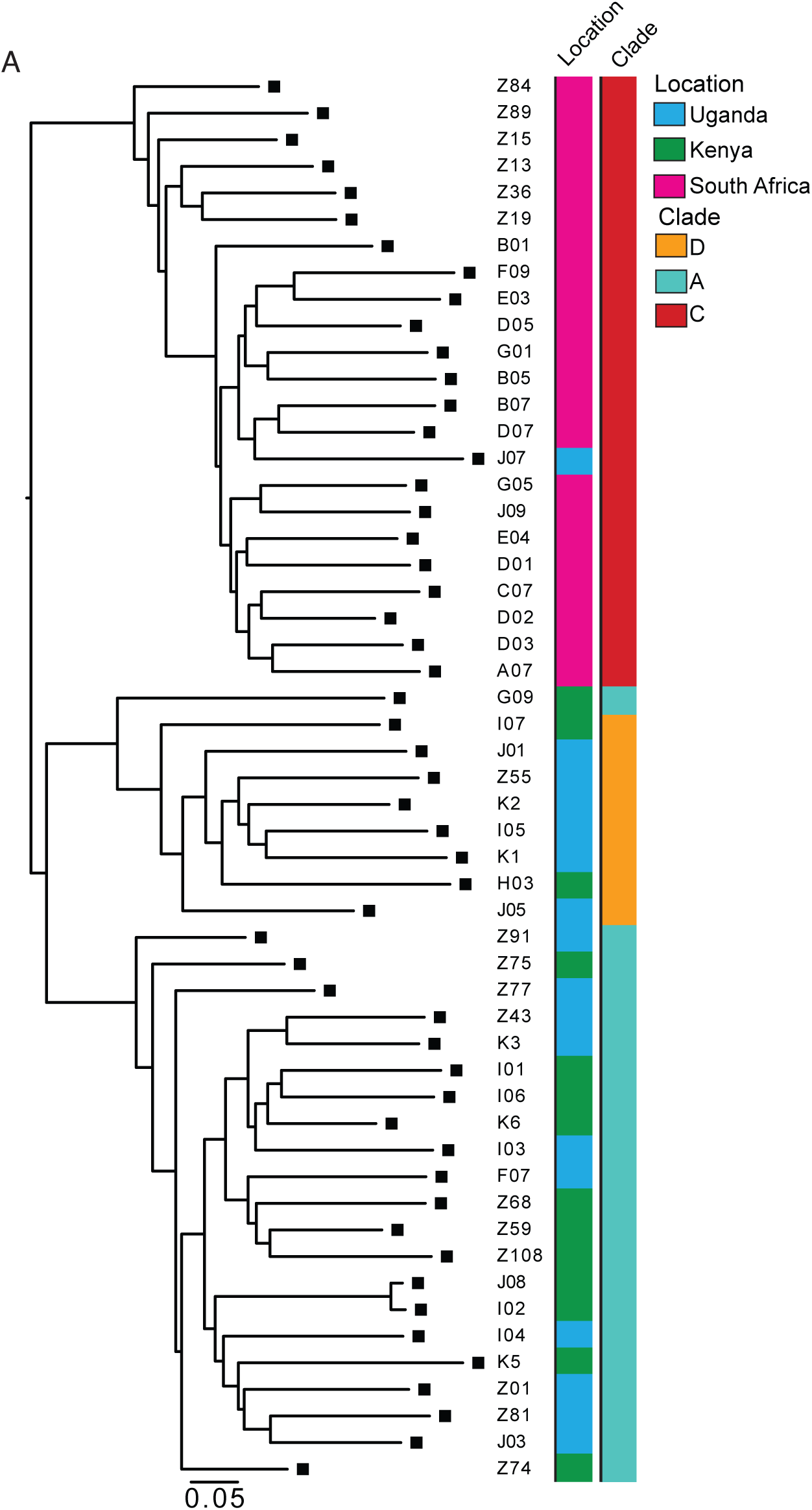
(a). Maximum likelihood phylogenetic tree of *env* sequences of viruses isolated from outgrowth cultures by SGA. The first bar to the right of the phylogenetic tree represents the country origin of sample collection, the second bar represents the HIV-1 clade of every sample. The heatmap represents the IC_80_ titer (µg/ml) for all samples for each bNAb tested.

The following bNAbs were tested for neutralizing activity against the PBMC isolates in TZM-bl assays: 3BNC117-LS, VRC01, VRC07-523LS, and 1-18, all of which are CD4 binding site-specific (CD4bs), (Scheid et al., 2011; Wu et al., 2010; Rudicell et al., 2014; Schommers et al., 2020); 10-1074-LS, and BG18, which target base of the V3 loop and surrounding glycans (Mouquet et al., 2012; Freund et al., 2017); and PGDM1400 and CAP256-VRC25.26, which are specific for the V2 loop (Sok et al., 2014; Doria-Rose et al., 2016). We also tested the combination of 3BNC117-LS and 10-1074-LS, which is currently in clinical development (Mendoza et al., 2018; Bar-On et al., 2018; Cohen et al., 2019)

(<u>clinicaltrials.gov/ct2/show/NCT03254277</u>, <u>clinicaltrials.gov/ct2/show/NCT03554408</u>, <u>clinicaltrials.gov/ct2/show/NCT04250636</u>, and <u>clinicaltrials.gov/ct2/show/NCT04173819</u>). The geometric mean IC_50_ for VRC01, which is now being tested in two large efficacy prevention trials (<u>clinicaltrials.gov/ct2/show/NCT02568215</u> and <u>clinicaltrials.gov/ct2/show/NCT02716675</u>), was 7.01 µg/ml for all viral isolates (Fig. 2a, supplementary table 2). Only 57% of the viruses tested were sensitive to VRC01 at concentrations below 10 µg/ml (Fig. 2a, b, c, d, Table 1, Supp table 2). Other CD4bs antibodies were substantially more potent than VRC01, including VRC07-523-LS and 1-18, with geometric mean IC_50_s of 1.14 µg/ml and 1.27 µg/ml, respectively. These two antibodies alone covered 92% and 87% of the viruses tested at concentrations below 10 µg/ml (Fig. 2a, b, c, d, Table 1, Supp table 2). 10-1074-LS and BG18, which target the base of the V3 loop, demonstrated geometric mean IC_50_s of 2.55 µg/ml and 2.64 µg/ml, respectively (Fig. 2a, b, c, d, Table 1, Supp table 2). PGDM1400 and CAP256-VRC25.26, which target the V2 loop, demonstrated geometric mean IC_50_s of 3.38 µg/ml and 0.16 µg/ml, respectively. However, 10-1074-LS, BG18, PGDM1400 and CAP256-VRC25.26 covered only 52%, 49%, 46% and 47% of the viruses, respectively, at concentrations below 10 µg/ml (Fig. 2a, b, c, d, Table 1, Supp table 2). The combination of 3BNC117-LS and 10-1074-LS performed better than any single antibody alone in terms of potency. The geometric mean IC_50_ for the combination was 0.65 µg/ml, and 84% of the viruses were sensitive at concentrations below 10 µg/ml (Fig. 2a, b, c, d, Table 1, Supp table 2).

**Figure 2.**
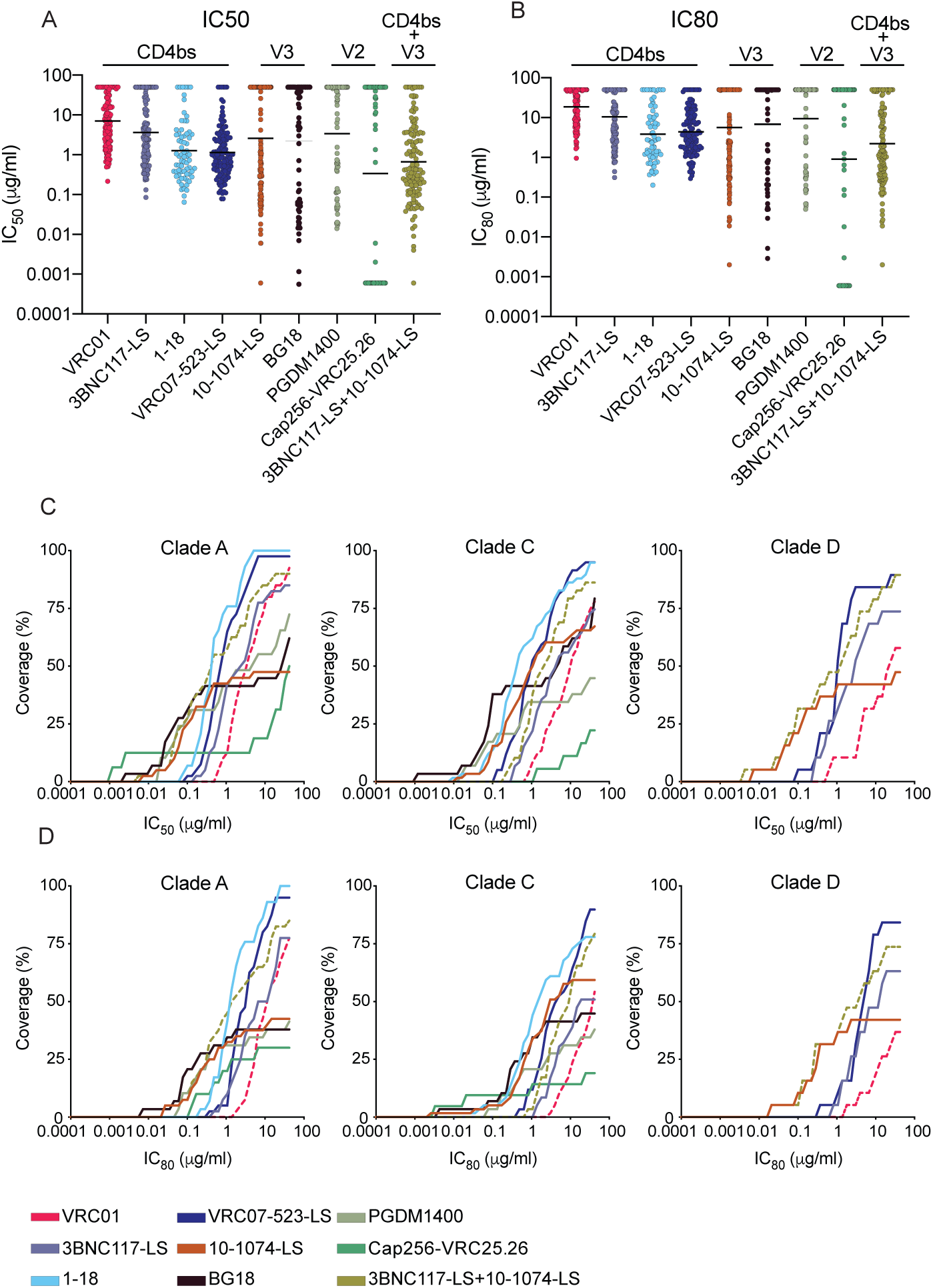
(a) Dot plot representing the IC_50_ titer (µg/ml) of unique PBMC-derived viruses for each bNAb tested. (b) Dot plot representing the IC_80_ titer (µg/ml) of unique PBMC-derived viruses for each bNAb tested. Each dot represents a single virus. Black bars represent geometric mean IC_50/80_ titers. (c and d) Coverage curves. For each antibody, the graph shows the percentage of viruses neutralized in the TZM-bl assay at a given (c) IC_50_ titer (µg/ml) or (d) IC_80_ titer for the PBMC-derived primary isolates for HIV-1 clades A, C and D. All bNAbs are represented with the same color scheme as in (a). The pink dotted line represents VRC01 and the olive green represents the combination of 3BNC117-LS and 10-1074-LS.

To determine whether the sensitivity of the primary African isolates to bNAbs differs from standard pseudovirus panels, we compared the data obtained from the outgrowth cultures with well characterized clade A, C, and D pseudoviruses (Fig. 3a, b and Supp. Table 3 and 4). All of the bNAbs tested were more potent and showed increased breadth against the pseudoviruses compared to the primary isolates. The difference between pseudovirus and primary isolates varied between antibodies. For example, CD4 binding site-specific bNAbs showed an average decrease in potency of 20-13- and 27-fold for primary isolates from clades A, C, and D, respectively. Moreover, these antibodies neutralized an average of 4.2%, 13.5%, and 28.1% fewer clade A, C, and D primary isolates, respectively, then when tested against primary isolates compared to pseudoviruses at concentrations below 10 µg/ml.

**Figure 3.**
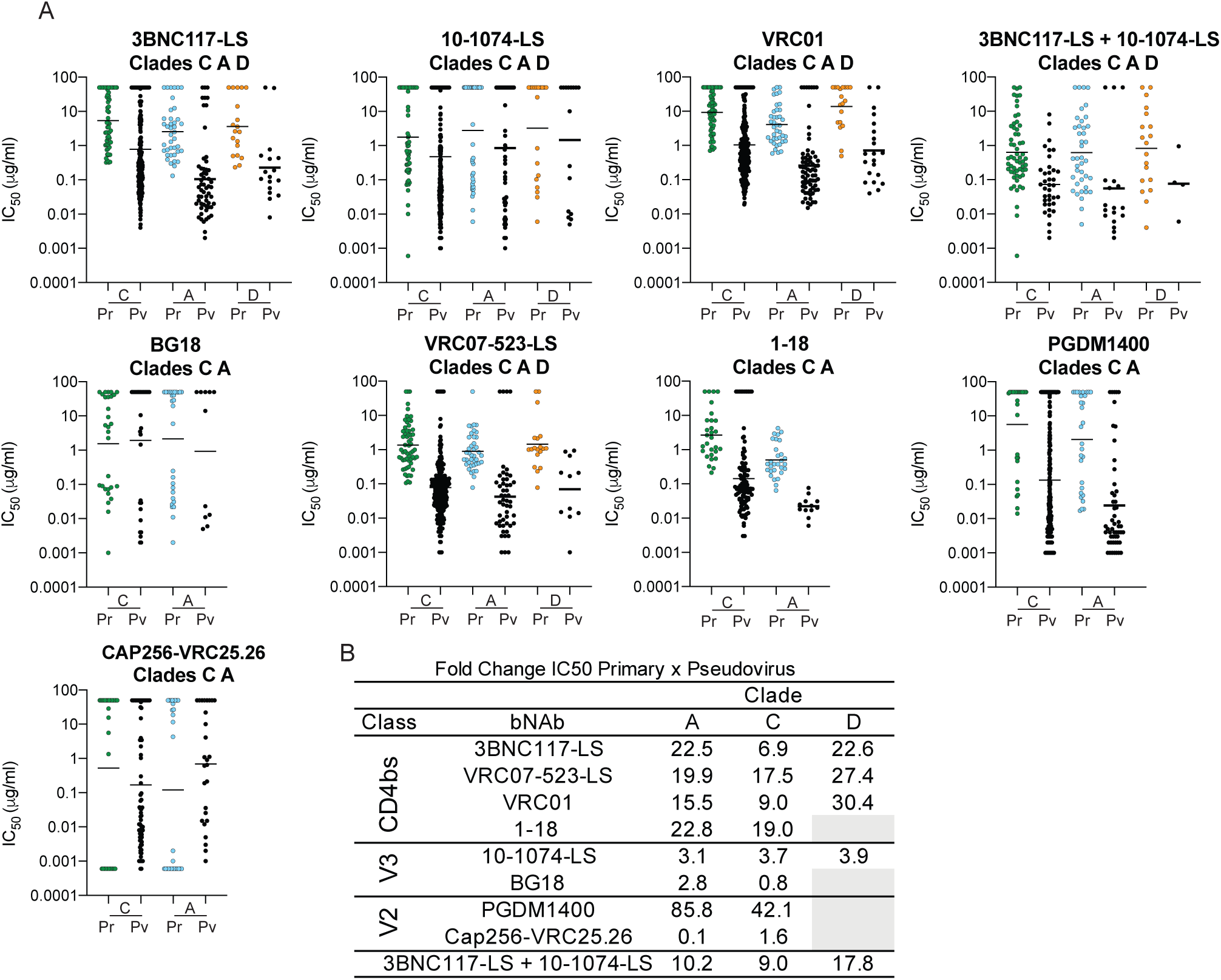
(a). IC_50_ titer (µg/ml) of unique PBMC-derived viruses (Pr) shown in color, and corresponding clade pseudovirus panel in black (data obtained from the Antibody database software (West et al., 2013); CATNAP (Yoon et al., 2015);, (Freund et al., 2017); (Schommers et al., 2020) and (Kong et al., 2015) (Pv). The viruses are organized by HIV-1 clade. Each dot represents a single virus. Black bars represent geometric mean IC_50_ titers. (b) Table describing the fold change between the geometric mean IC_50_ titer (µg/ml) between PBMC-derived viruses and the corresponding pseudoviruses panel.

The difference in potency and breadth between pseudoviruses and primary isolates was less dramatic for bNAbs targeting the V3 loop. On average, there was only a 3-fold difference in IC_50_ between primary isolates and pseudovirus panels for clades A, C, and D. V3-loop antibodies also retained most of their breadth when considering the numbers of strains reaching IC_50_s at concentrations below 10 µg/ml (Fig. 3a, b and Supp. Table 3 and 4).

The two V2-loop bNAbs were unusual in that they were very different in their relative potencies against primary and pseudotyped Clade A and C viruses. Whereas CAP256-VRC25.26 showed only a small difference in activity, PGDM1400 was 85- and 42-fold less active against primary clade A and C viruses than pseudotyped viruses, respectively (Fig. 3a, b and Supp. Table 3 and 4). These antibodies neutralized 24% and 32% fewer clade A and C primary isolates, respectively, than pseudoviruses at concentrations below 10 µg/ml. Finally, the 3BNC117-LS/10-1074-LS combination was on average 12-fold less active against the primary isolates than pseudoviruses, and showed no decrease in breadth for clade A, but 13.5%, and 26.5% decrease in breadth when considering the numbers of strains reaching IC_50_s at concentrations below 10 µg/ml for clades C and D respectively (Fig. 3a, b and Supp. Table 3 and 4).

To determine whether the differences between primary isolates and pseudovirus panels was attributed to sequence differences between the viruses being tested, we cloned HIV-1 env genes from 13 different primary cultures, expressed them as pseudotyped viruses and tested them against a panel of 5 bNAbs in the TZM-bl neutralization assay. PBMC-derived viruses and the matched pseudoviruses IC_50_s showed similar fold differences than those found between primary isolates and pseudovirus panels for all bNAbs tested (Fig. 4). The data suggest that there are significant differences in bNAb potency and breadth between primary clade A, C and D isolates and pseudotyped viruses, and that the magnitude of these differences is bNAb specific.

**Figure 4.**
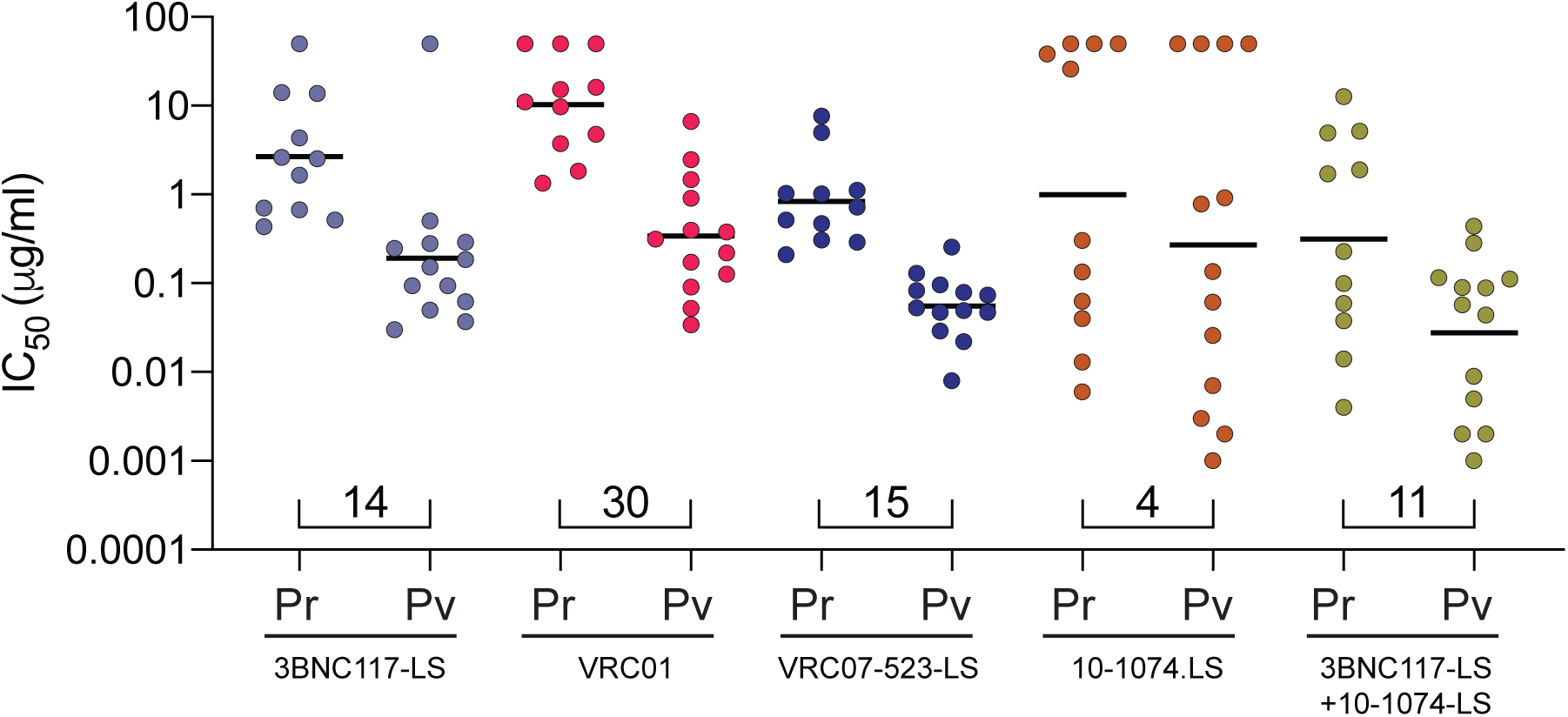
(a) IC_50_ titer (µg/ml) of unique PBMC-derived clonal viruses (Pr) and corresponding pseudoviruses (Pv) for each antibody. Each dot represents a single virus. Black bars represent geometric mean IC_50_ titers. Numbers under the dots indicate the fold change in geometric mean IC_50_ titers between the 2 groups.

## Discussion

We measured the neutralization profile of nine bNAbs currently in clinical development on 126 primary isolates obtained from PBMC cultures from individuals infected with HIV-1 clades A, C, and D. VRC01, the most advanced clinical candidate, is nearly 15 times less active against primary isolates than pseudotyped viruses. Similar results were obtained with other CD4 binding site antibodies. In contrast, the two V2-directed antibodies tested varied widely in their ability to neutralize pseudotyped viruses and primary isolates. Thus, the results obtained with pseudotyped virus panels cannot be translated directly to bNAb activity on primary isolates.

Our results extend earlier work with less potent antibodies (Louder et al., 2005; Mann et al., 2009; Provine et al., 2012), and with bNAbs against clade B viruses (Cohen et al., 2018b) to clades A, C and D. In all cases, primary isolates were less sensitive to bNAbs than pseudotyped viruses. However, the relative reduction in activity differed between antibodies that target different epitopes on the envelope spike, with V3-loop bNAbs 10-1074-LS and BG18 being least affected and PGDM1400 the most affected. In addition, the magnitude of the differences varies among viral clades. Combinations of bNAbs, as exemplified by 3BNC117-LS and 10-1074-LS, are advantageous in this respect, as also suggested by *in vitro* and in silico analysis using Env-pseudotyped panels (Zwick et al., 2001; Goo et al., 2012; Doria-Rose et al., 2012; Wagh et al., 2016, 2018; Kong et al., 2015).

A number of non-mutually exclusive hypotheses have been suggested to explain the enhanced susceptibility of 293T-derived pseudotyped viruses to neutralization by bNAbs. For example, sensitivity to neutralization could be dependent on the number of envelope protein spikes, with fewer spikes bound on the surface of 293T-derived pseudotyped viruses than PBMC derived primary isolates (Louder et al., 2005; Provine et al., 2012). Another possibility involves differential glycosylation by different packaging cell types. bNAbs frequently target glycan dependent epitopes, therefore the differential glycosylation profile of the envelope spike produced in different cell types could also alter their neutralization profile (Duenas-Decamp et al., 2008; Utachee et al., 2010). However, V3-loop bNAbs and CAP256-VRC25.26 that target highly glycan dependent epitopes were the least affected. Similarly, PG9, a V2 peptide-glycan specific bNAb (Walker et al., 2009; Doores and Burton, 2010), showed only small changes in its neutralization profile between clade B pseudotyped viruses and PBMC-derived viruses (Provine et al., 2012).

Clinical trials testing bNAbs for HIV-1 prevention are now being conducted in Africa and other parts of the world. The largest of these trials is testing VRC01 at several sites in Africa (Botswana, Kenya, Malawi, Mozambique, South Africa, Tanzania and Zimbabwe), where the majority of the HIV-1 infections are caused by clades A, C, and D viruses (Bbosa et al., 2019). Although the results of those trials are not yet known, data are available from smaller trials where bNAbs were administered to individuals undergoing analytical treatment interruption (ATI). In the absence of antiretroviral therapy, nearly all participants experience viral rebound in 2-3 weeks, and it is believed that recrudescence of viremia is due to reactivation of HIV-1 from latently infected CD4^+^ T cells (Sengupta and Siliciano, 2018). Single antibodies were able to decrease viremia levels or delay the return of quantifiable viremia, but their ability to do so correlated with their neutralizing activity against primary isolates and not pseudotyped viruses (Caskey et al., 2015; Scheid et al., 2016; Caskey et al., 2017; Cohen et al., 2018a; Stephenson et al., 2019; G. Chen et al, 2019). For example, VRC01 had little measurable effect on delaying HIV-1 rebound when administered during ATI (Bar et al., 2016; Crowell et al., 2019). In contrast, antibody combinations maintain suppression of viremia during ATI in individuals harboring bNAb sensitive viruses for as long as antibody concentrations remain above 10 µg/ml (Mendoza et al., 2018). Should the clinical outcomes in the ongoing VRC01 prevention trials track with bNAb activity against primary isolates as opposed to pseudotyped virus panels, there could be up to a 15-fold difference between the predicted and observed outcomes of the trial.

Nevertheless, by analogy with the ATI trials, if the AMP trials demonstrate even a smaller than projected effect with VRC01, it provides a proof-of-concept that passive immunization can prevent sexual transmission of sensitive HIV-1 strains and indicate that combinations should be highly effective.

## Methods

### Samples

The study was conducted under the approval of The Rockefeller University Institutional Review Board. Samples were collected during the course of three studies in sub-Saharan Africa: (1) The Partners in Prevention HSV/HIV Transmission Study: between November 2004 and April 2007, 3408 HIV serodiscordant heterosexual couples were enrolled from 14 study sites in sub-Saharan Africa into this phase III clinical trial evaluating the efficacy of HSV-2 suppressive therapy (acyclovir 400 mg orally twice-daily versus matching placebo) provided to persons infected with both HIV-1 and HSV-2 who had CD4 count ≥250 at enrollment to prevent HIV transmission to their HIV-uninfected heterosexual partner (Celum et al., 2010); (2) Couples Observational Study: A total of 485 HIV serodiscordant heterosexual couples were recruited at two of the same sites as the Partners in Prevention HSV-2/HIV Transmission Study (Kampala, Uganda and Soweto, S. Africa) for a prospective, observational study of biologic correlates of HIV protection; there was no HSV-2 co-infection or CD4 count enrollment requirement (Lingappa et al., 2011); (3) Partners PrEP Study: this was randomized, phase III clinical trial of antiretroviral pre-exposure chemoprophylaxis (300 mg tenofovir once daily versus 300 mg Tenofovir/200 mg emtricitabine once daily versus matching placebo) conducted at nine sites in Kenya and Uganda (Baeten et al., 2012).

### CD4^+^ T cell outgrowth culture

Bulk outgrowth cultures were performed as previously described (Cohen et al., 2018a). Briefly, PBMCs were obtained from HIV-1-infected individuals, and CD4^+^ T lymphocytes were isolated by negative selection with magnetic beads (Miltenyi). A total of 2×10^6^ CD4^+^ T lymphocytes were activated using anti CD3/CD2/CD28 beads (Miltenyi) and cultured in the presence of 100 U/mL IL-2 (Peprotech) at 37 °C and 5% CO2. CD4^+^ T lymphocytes were co-cultured with irradiated heterologous health donor PBMCs (1×10^6^). After 24 hours of activation, 1×10^5^ Molt 4 CCR5 cells were added. The medium was replaced twice a week, and the presence of p24 in the culture supernatant was quantified by the Lenti-X p24 Rapid Titer kit (Clontech) after 7, 14 and 21 days of culture. The infectivity of viral cultures was confirmed by a 50% tissue culture infective dose assay with TZM-bl cells (Li et al., 2005)

### Neutralization assays

TZM-bl cell neutralization assays were performed as previously described (Montefiori, 2005; Li et al., 2005). Neutralization assays were conducted in laboratories meeting Good Clinical Laboratory Practice quality assurance criteria. All bulk outgrowth culture primary isolates were tested against 3BNC117-LS, VRC01, 10-1074-LS, VRC07-523-LS and the combination of 3BNC117-LS and 10-1074-LS. Sixty nine were also tested against PGDM1400 (provided by Dennis Burton, Scripps Research Institute), BG18, 1-18 (provided by Florian Klein, University of Cologne) and CAP256-VRC25.36 (provided by John Mascola, NIH Vaccine Research Center). The maximum antibody concentration tested was 50 μg/ml.

### Virus sequence analysis

HIV *env* sequences from p24-positive supernatants were obtained and analyzed as previously described (Lorenzi et al., 2016). Sequences derived from each bulk culture with double peaks (cutoff consensus identity for any residue <75%), stop codons, or shorter than the expected envelope size were omitted from downstream analysis. Phylogenetic analysis was performed by generating nucleotide alignments using MAFFT (Katoh and Toh, 2010) and posterior phylogenetic trees using PhyML v3.1 (Guindon et al., 2010), using the GTR model with 1000 bootstraps. Clade determination was performed using the NCBI subtyping tool (http://www.ncbi.nlm.nih.gov/projects/genotyping/formpage.cgi). For samples not sequenced in this study the clade was determined by sequencing a 514 base pair (bp) region of the *env* gene (C2-V3-C3 region) from plasma samples as previously described (Celum et al., 2010; Lingappa et al., 2011; Baeten et al., 2012).

### Pseudotyped virus production

Pseudotyped virus production was performed as previously described (Kirchherr et al., 2007). The cytomegalovirus (CMV) promoter was amplified by PCR from the pcDNA 3.1D/V5-His-TOPO plasmid (Life Technologies) with forward primer 5’-AGTAATCAATTACGGGGTCATTAGTTCAT-3’ and reverse primer 5’ CATAGGAGATGCCTAAGCCGGTGGAGCTCTGCTTATATAGACCTC-3’. A 1μl volume of the first-round PCR product from each individual *env* gene obtained from bulk cultures was amplified with primers 5’CACCGGCTTAGGCATCTCCTATGGCAGGAAGAA-3’ and 5’ACTTTTTGACCACTTGCCACCCAT-3’. PCR products were purified with the Macherey-Nagel Gel and PCR purification kit. The CMV promoter amplicon was fused to individual *env* genes by overlap PCR with 10 ng of *env* and 0.5 ng of CMV with forward primer 5’-AGTAATCAATTACGGGGTCATTAGTTCAT-3’ and reverse primer 5’-ACTTTTTGACCACTTGCCACCCAT-3’. Resulting amplicons were analyzed by gel electrophoresis, purified with the Macherey-Nagel Gel and PCR purification kit, and co-transfected with pSG3Δenv backbone vector (NIH AIDS Reagent Program) into HEK293T cells to produce pseudoviruses as previously described (Kirchherr et al., 2007).

## Acknowledgements

We thank the study participants who devoted time to our research and all members of the Nussenzweig lab for helpful discussions. This work was supported in part by Bill and Melinda Gates Foundation Collaboration for AIDS Vaccine Discovery (CAVD) grants OPP1146996 (M.S.S.), OPP1092074, and OPP1168933 (M.C.N.); the NIH Center for HIV/AIDS Vaccine Immunology and Immunogen Discovery (CHAVI-ID) 1UM1 AI100663-06 (M.C.N.); M.C.N. is a Howard Hughes Medical Investigator.

## Supplemental Material

**Supplementary Table 1**. Demographic characteristics of individual participants. M – Male, F – Female.

**Supplementary Table 2**. Neutralization profile of PBMC derived samples and the respective IC50s and IC80 for all tested bNAbs.

**Supplementary Table 3**. Potency and breadth of all tested bNAb against HIV isolates from PBMC derived samples or pseudovirus for each clade.

**Supplementary Table 4**. Total number of tested samples for individual bNAb and clade. PBMC – PBMC derived primary samples, PV – Pseudovirus.

**Supplementary table 1.** Demographic characteristics of tested individuals.

| Country | Total tested | All Tested |  |  |  |  |  |  |  | Positive Cultures |  |  |  |  |  |  |  |
| --- | --- | --- | --- | --- | --- | --- | --- | --- | --- | --- | --- | --- | --- | --- | --- | --- | --- |
|  |  | Clade |  |  |  | Av.Age |  | Sex |  | Clade |  |  |  | Av.Age |  | Sex |  |
|  |  | A | C | D | nd | F | M | F | M | A | C | D | nd | F | M | F | M |
| South Africa | 84 | 1 | 58 | 0 | 25 | 31.2 | 37.0 | 57 | 27 | 0 | 56 | 0 |  | 31 | 36.9 | 41 | 15 |
| Kenya | 66 | 27 | 4 | 11 | 24 | 28.8 | 37.4 | 44 | 22 | 21 | 1 | 1 | 5 | 28 | 35.2 | 18 | 10 |
| Uganda | 68 | 35 | 3 | 23 | 7 | 28.3 | 32.1 | 36 | 32 | 25 | 2 | 14 | 1 | 28 | 33.1 | 22 | 20 |

F - Female, M - Male, nd - not determined

**Supplementary table 2.**
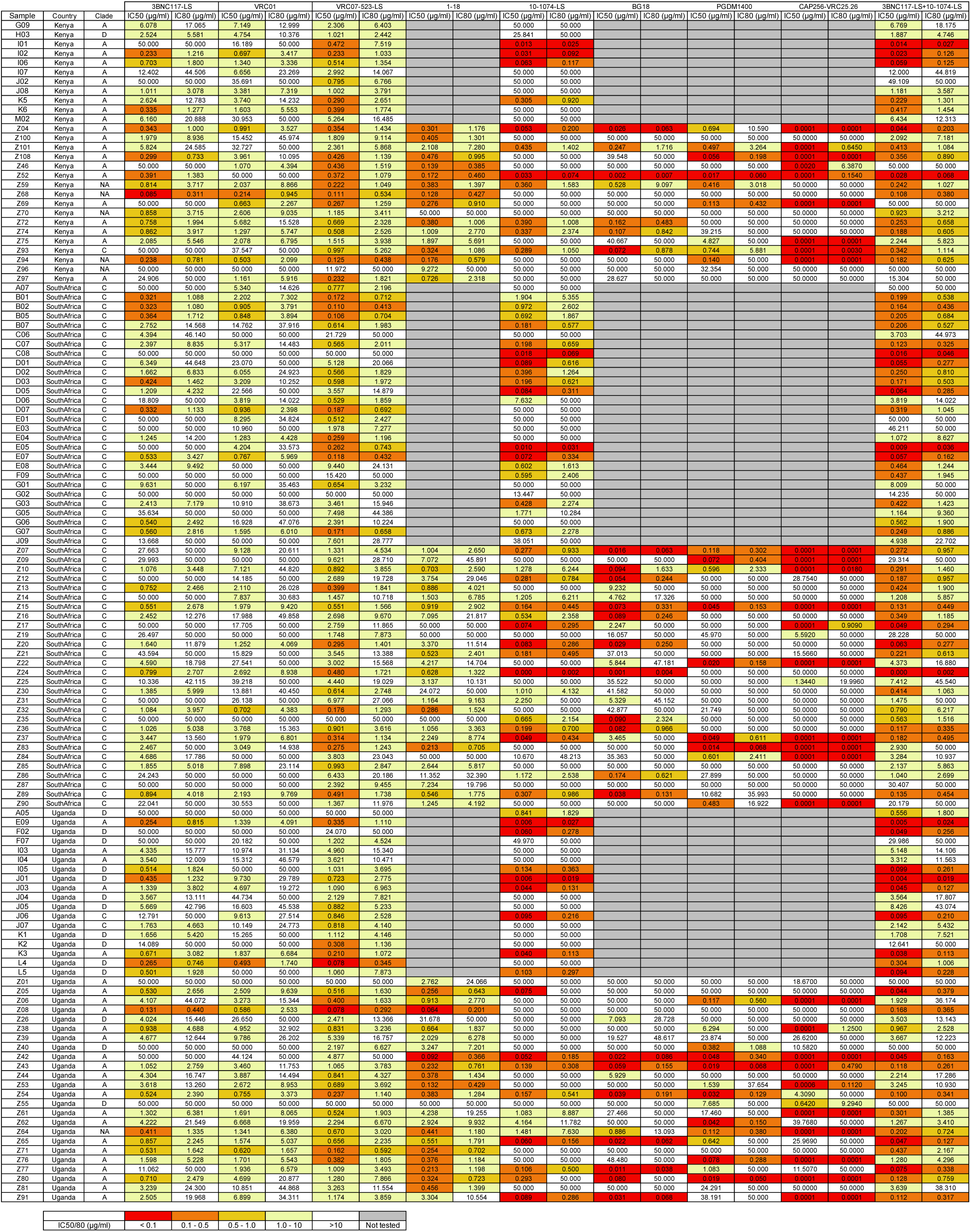
Neutralization profile of PBMC derived samples and the respective IC_50_s and IC_80_s for all tested bNAbs

**Supplementary table 3.**
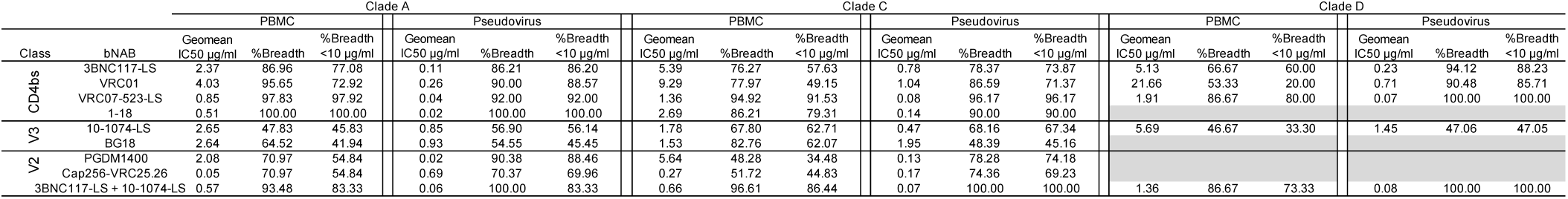
Potency and breadth of bNAbs tested against PBMC derived isolates and pseudovirus panels defined by HIV clade.

**Supplementary table 4.** Total number of tested samples for individual bNAb and clade. Pr – PBMC derived primary samples, Pv – Pseudovirus

| Clade |  | A |  | C |  | D |  |
| --- | --- | --- | --- | --- | --- | --- | --- |
| Class | bNAb | Pr | PV | Pr | PV | Pr | PV |
| CD4bs | 3BNC117-LS | 46 | 58 | 59 | 245 | 15 | 17 |
|  | VRC01 | 46 | 50 | 59 | 235 | 15 | 11 |
|  | VRC07-523-LS | 46 | 70 | 59 | 276 | 15 | 21 |
|  | 1-18 | 31 | 12 | 29 | 100 |  |  |
| V3 | 10-1074-LS | 46 | 57 | 59 | 245 | 15 | 17 |
|  | BG18 | 31 | 11 | 29 | 31 |  |  |
| V2 | PGDM1400 | 31 | 52 | 29 | 244 |  |  |
|  | Cap256-VRC25.26 | 31 | 27 | 29 | 78 |  |  |
| 3BNC117-LS + 10-1074-LS |  | 46 | 18 | 59 | 37 | 15 | 4 |

